# Rattractor - Instant guidance of a rat into a virtual cage using a deep brain stimulation

**DOI:** 10.1101/2022.10.07.511260

**Authors:** Naoki Sudo, Sei-etsu Fujiwara, Takashi Isoyama, Osamu Fukayama

## Abstract

We developed “Rattractor” (rat attractor), a system to apply electrical stimuli to the deep brain of a rat as it stays in a specified region or a virtual cage to demonstrate an instant electrophysiological feedback controller for animals.

Two wire electrodes were implanted in the brains of nine rats. The electrodes targeted the medial forebrain bundle (MFB), which is a part of the reward system in the deep brain. Following the recovery period, the rats were placed in a plain field where they could move freely, but wired to a stimulation circuit. An image sensor installed over the field detected the subject’s position, which triggered the stimulator such that the rat remained within the virtual cage. We conducted a behavioral experiment to evaluate the sojourn ratio of rats residing in the region. Thereafter, a histological analysis of the rat brain was performed to confirm the position of the stimulation sites in the brain.

Seven rats survived the surgery and the recovery period without technical failures such as connector breaks. We observed that three of them tended to stay in the virtual cage during stimulation, and this effect was maintained for two weeks. Histological analysis revealed that the electrode tips were correctly placed in the MFB region of the rats. The other four subjects showed no apparent preference for the virtual cage. In these rats, we did not find electrode tips in the MFB, or could not determine their positions.

Almost half of the rats tended to remain inside the virtual cage when position-related reward stimuli were triggered in the MFB region. Notably, our system did not require previous training or sequential interventions to affect the behavioral preferences of subjects. This process is similar to the situation in which sheep are chased by a shepherd dog in the desired direction.

## Introduction

Electrical stimulation in living animals can induce virtual sensations and muscular contractions, resulting in behavioral alternations. For example, Lopez et al. [1] developed various virtual reality games in which players experienced somatosensory impacts on their limbs by electrically triggering muscle contraction in virtual environments such as a boxing ring or a soccer field. Another example of this technology is “Cruise Control” [2] which providing automated muscle stimuli, allowed a person to avoid obstacles on a predefined pathway with their eyes closed. Moreover, it has been applied to posture training when pitching a bowling ball by inducing muscular contraction in the limb [3].

Electrical stimulation of muscles, also called functional electrical stimulation (FES), has long been used in clinical applications for patients with paralysis or spinal cord injuries to manipulate their disabled limbs to grasp objects, stand up, and even step forward [4]. Recently, a method for bypassing injured nerve fibers and transmitting efferent voluntary motor commands in the patient’s own muscles has been proposed [5].

In animal models, more direct behavioral interventions have been demonstrated. *Roboroach* [7–9], for example, applied stimuli to the hemi-lateral antennae and manipulated cockroach locomotion in the opposite direction.

The behavior was manipulated by an experimenter who dispatched sequential instructions according to the circumstances, similar to a radio-controlled vehicle.

In contrast, *Ravigation* or a rat navigation system [6] affects rodent behavior through a rewarding stimulus in the medial frontal bundle (MFB) region. The subject received stimuli in the MFB, which is a part of the reward system in the brain. The subject was rewarded after following instructions to make a directional turn indicated by an electrical stimulus in the sensory cortex (SCx). However, this system also required sequential instructions and ex ante training periods for the subject to associate the stimuli in the SCx and MFB. Training is not only a burden for both experimenters and subjects but also increases the possibility of unexpected interruptions such as device defects and animal fatigue.

To avoid these problems, we developed an automated reward stimulator in which the rat position triggered stimuli. This affects animal behavior without training, and could be useful for industrial applications based on animal behavior, such as mine exploration in minefield barrier fences. We developed a system called “Rattractor” as a proof-of-concept, in which the reward system in the rat brain was stimulated when the subject stayed in a specific circular region simulating a virtual cage. While a previous study on the system roughly demonstrated this concept [10], this study goes one step further by focusing on the quantitative evaluation of the system.

## Materials & methods

### Methodology

This section describes the general approach of the Rattractor system including animals preparation.

#### System Design

##### Devices

The Rattractor system is essentially an electrical feedback stimulator. It stimulates the rat brain using electrical stimuli triggered by an optically observed rat position. To achieve this functionality, we placed a high-speed, low-latency camera (Basler acA800-510uc) over a plain field (1.4 m × 0.9 m) where a rat can freely explore, wander, and rest (Fig 1). The top surface of the field was covered with a black elastic sheet, and three edges were limited by walls also covered with elastic sheets, while the other edge faced a cliff from which rats rarely jumped out.

**Fig 1.**
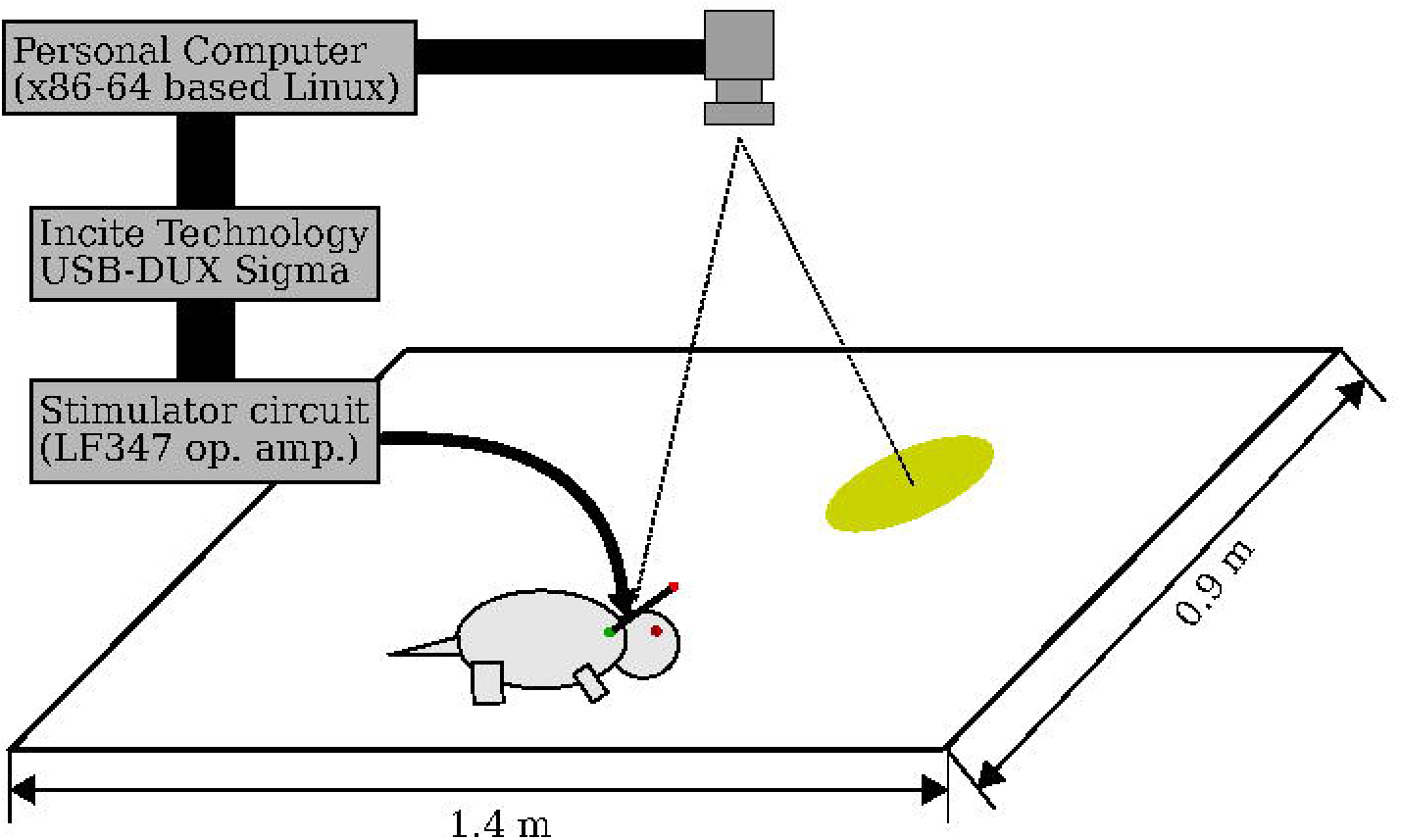
Appearance of the Rattractor system. The rat was stimulated in the MFB as it entered the virtual cage indicated with a circle.

The camera, which had XGA (1024 × 768) resolution, covered the entire field including its edges, and was connected to a desktop computer (Intel Core i5 CPU; 8 GB memory; operated with Linux) using a USB 3.0 cable. After detecting the rat’s position, the computer produced stimulation triggers via an analog output interface (Incite Technology, USB-DUX Sigma).

We attached a current source circuit (Fig 2a) to the analog output connector, which consisted of a voltage differentiation circuit and a constant current supplier circuit, using an LF347 operational amplifier package to inject charges into a living animal. The circuit accepts two monopolar voltage inputs *V*_1_ and *V*_2_ between 0–2.5 V from the analog output board to generate currents *I* = (*V*_1_ – *V*_2_)/10^4^ between −2.5 and +2.5 mA. It also included a shunt resistor to monitor a voltage proportional to the current.

**Fig 2.**
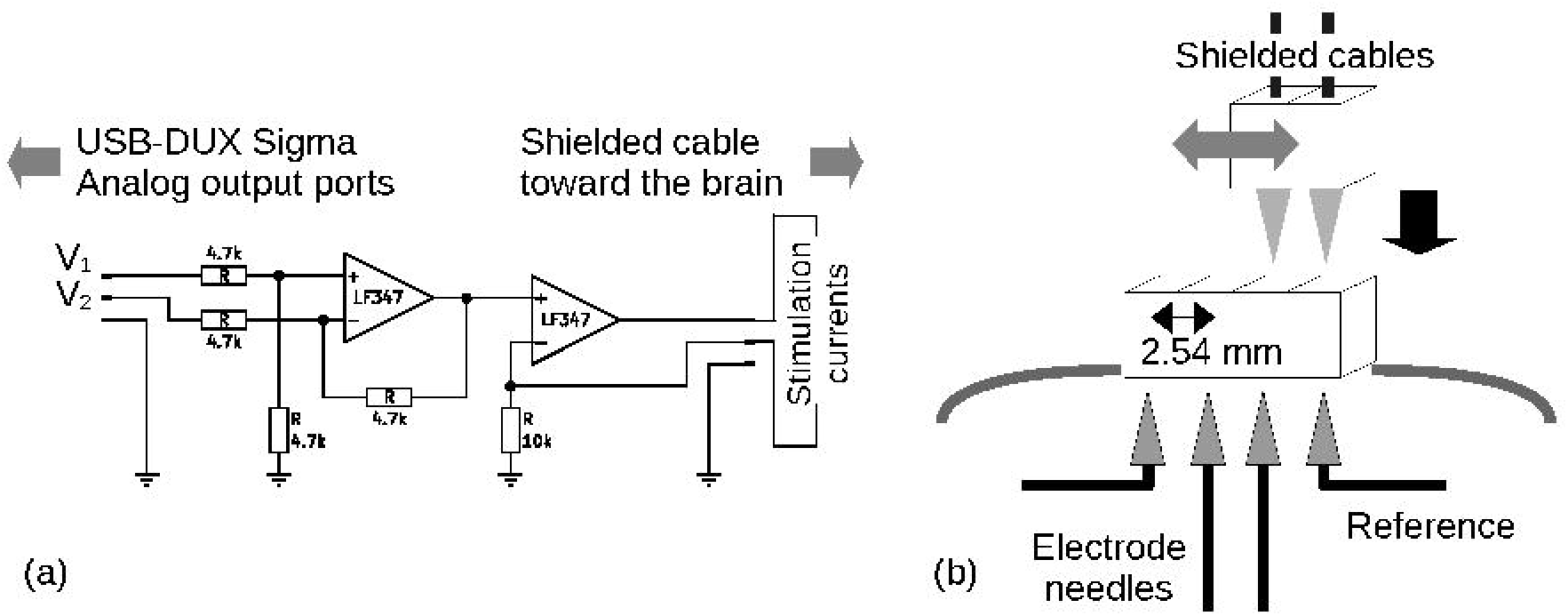
(a) Stimulator circuit to generate bipolar current stimuli. (b) Electrode unit with two electrode needles and two references.

Currents were then applied to the brain through a stimulation interface unit placed on the skull. The unit consisted of two needles and a conventional 0.1 inches (2.54 mm) pitched connector with four lines (Fig 2b). Each needle was clipped from a commercially available tungsten neural macro electrode (LF-301G; MicroProbes).We removed the polyamide coating around its end using a paper grinder and swaged the metal contact pin at the end. The needles were then inserted into the second and third terminals at an interval of 0.1 inches (2.54 mm) between the two needles. The first and fourth terminals were used to hold the reference lines. Finally, the entire structure was covered and fixed using thermoplastic adhesive.

The unit was wired to the stimulator circuit using shielded cables during behavioral experiments. Two green and red light-emitting diodes (LEDs) connected by a battery (CR2032) were placed at the end of the cables, which facilitated tracking of the position and direction of the rat’s head.

##### Controller

A controller was implemented on the computer to dispatch stimulation triggers according to the position of the rat. Image frames were grabbed every 30 – 50 ms. Then, green and red color filters and binarization were applied to the image, and their spatial moments were calculated to obtain ***r***_Green_ and ***r***_Red_, the coordinates of the red and green LEDs, respectively.

The rat position was calculated as follows: ***r***_Rat_ = (***r***_Green_ + ***r***_Red_)/2, which triggered the stimulus as long as the rat remained within the virtual cage. That is, stimulation was triggered as |***r***_Rat_ – ***r***_Cage_| < 0.1 m, where ***r***_Cage_ was the position of the virtual cage. To avoid excessive brain stimulation, the trigger frequency was limited to a maximum of 5 Hz.

Each trigger generated a waveform consisting of ten repeating bipolar squares at an amplitude of 80 *μ*V and a duration of 5 ms (Fig 3).

**Fig 3.**
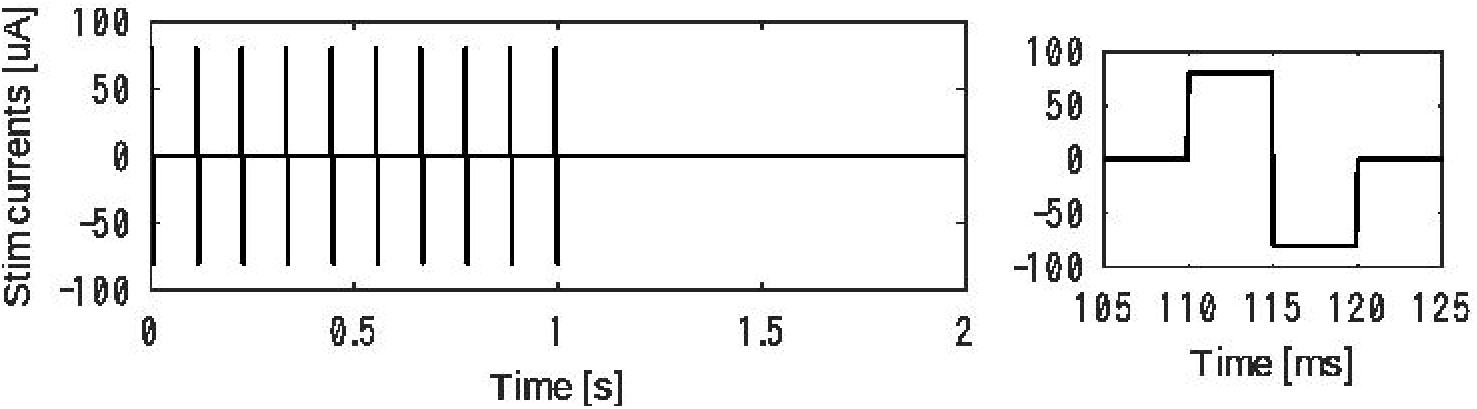
A set of 10 square pulses were generated with our circuit to apply bipolar current stimuli (left); temporal closeup around the first pulse (right).

#### Animals preparation

Nine male Wistar rats were prepared for the experiment. Animals were bred by an animal provider (Tokyo Laboratory Animals Science Co., Ltd.) in a clean/SPF-graded environment until delivery to our facility. At this point, they weighed 250 g and were approximately 10-weeks-old. They were able to drink water freely in the cages but were fed a restricted amount of solid food to maintain their weight until surgery, which was typically conducted 1–4 weeks after arrival.

Surgery to implant the electrode unit was conducted as follows: First, rats were anesthetized with isoflurane gas at 2.0 – 2.5 % and kept immobilized at 1.0 – 1.5 % until the end of the surgery. After applying Xylocaine jelly local anesthesia to the parietal skin, the rat was fixed to a stereotaxic brain surgery device. Then, we applied isodine to the skin followed by maskin hydrochloride for sterilization while waiting for the local anesthesia to be effective. Subsequently, we incised the skin and subcutaneous tissues using a surgical blade and drilled holes in the skull to implant both electrodes and anchors.

Two holes were drilled at 1.3 mm and 3.7 mm posterior, and 2.0 mm right lateral from the *bregma* skull joint using a hand drill with a 1.2 mm bit. These holes directed the needles into the MFB, which was found approximately along the body axis in deep regions of the brain. In addition, depending on the condition of the skull surface, six to eight holes were drilled to hold the anchor screws.

Finally, the electrode unit was placed over the two holes in the skull with a stereotaxic manipulator, and directed ventrally in the brain until it reached a depth of 8 mm from the brain surface.

### Experimental procedures

#### Behavioral Experiments

Behavioral experiments were conducted at 1 week (POW 1) and two weeks (POW 2) after surgery. All rats participated in POW 1 experiments, except for subject D, whose connectors accidentally detached during the recovery period. Based on their preference to stay in the virtual cage in the POW 1 session, three subjects (C, F, and I) proceeded to the POW 2 experiments.

The experiment began by wiring the rats to provide electrical stimuli to the brain. When the cables were connected to the two forward lines of the unit, the anterior MFB site was stimulated against the reference placed on the skull. Similarly, the posterior MFB site was stimulated against a reference placed on the skull by connecting the cables to the two rear lines of the unit.

The rat was then left to move freely in the field for 10 minutes for habituation. Most subjects exhibited exploratory behavior until they relaxed in the corner of the experimental field. Subsequently, we set a virtual cage region with our graphical user inferface (Fig 4), typically in front of the rat’s nose, so that the rat could enter the cage with only a single voluntary step.

**Fig 4.**
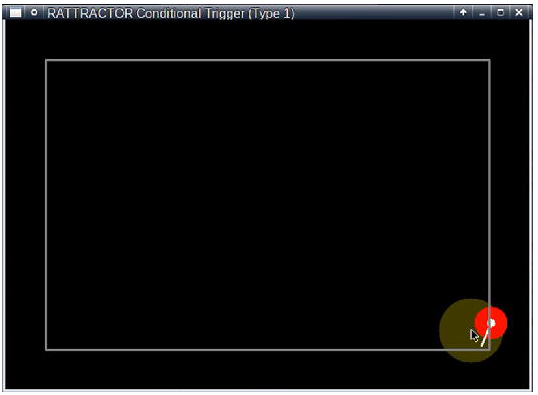
A graphical user interface (GUI) was developed to show the rat’s position (white circle) and heading direction (bar) as well as the stimulated region (i.e., virtual cage), drawn as a dark yellow circle and manually positioned by the experimenter using the mouse cursor. The red color around the rat position indicates that the subject is electrically stimulated at that time.

We prepared two conditions: the first *fixed* condition involved placing the virtual cage in the initial position for 10 minutes and the second *moving* condition involved moving the virtual cage straight from the initial position towards the far corner of the field. The *moving* condition was applied only if the stimuli in the *fixed* condition appeared to be effective.

#### Histology

After the behavioral experiments, the rats were deeply anesthetized with pentobarbital sodium, and electric lesions were burned with a 0.2 mA current to mark the electrode position. The brains were removed and stored in buffered formalin (10 %) for 7 days. Coronal sections (100 *μ*m) were cut using a microslicer (DTK-1000; DOSAKA EM, Japan) and stained with cresyl violet.

### Analysis

We analyzed both behavioral and histological results in two steps.

First, the rats were screened into two groups according to the stereotaxic coordinates of the electrode tips in the brain, histologically observed after sacrificing the rats. The subjects in whom the electrode tips were found in the MFB regions were sorted into the *positive* group, and their behavioral results on POW 1 and POW2 were analyzed. Other subjects in which we failed to find the electrode tips in the MFB were placed in the *negative* group, and their behaviors were only analyzed on POW1 as a reference session without reward stimuli.

We then defined a distance metric *d* = |***r***_Rat_ – ***r***_Cage_|, where ***r***_Rat_ and ***r***_Cage_ are the horizontal coordinates of the head position and center of the virtual cage, respectively. These metrics were calculated for every image frame at time stamp *k* (*k* = 1,…, *K*) captured by the camera for each subject (A, B,…, and H). In addition, another condition involving stimulated sites was evaluated. The stimulated sites were either the anterior, posterior, or null sites, where the currents were shunted.

The first 210 s (3 minutes and 30 seconds) of each session (*t_k_* = [0, 210]) were extracted to compare *d*(*t_k_*) values. The ratio of the frame count during which the rat stayed within the virtual cage was calculated as follows:

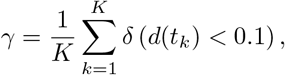

where *δ*(·) = 1 for a true condition, and *δ*(·) = 0 was calculated to compare the efficacy of attracting the rat between conditions, that is, subjects and stimulated sites.

In addition, on POW 2, the virtual cage tugging subject C was moved to an arbitrary location. During this session, the virtual cage moved from one corner to the opposite corner of the field and returned to the original center of the field.

### Remarks

The animal preparation and experimental procedures described in this section were conducted in an animal experiment facility in **** (anonymized).

## Results

### Screening

We inspected the rats’ appearance and behavior after the surgical procedure to determine whether the rats were suitable for subsequent experiments.

The behavioral screening focused on abnormal behaviors such as violent or fear responses during and after the electrical stimulation. We found no remarkable abnormal behavior instantly incudced by these stimuli. In addition, there were no persistent behavioral changes after stimulation;therefore, all the subjects participated in the following behavioral test sessions.

However, we removed several subjects from the behavioral analysis because of misplaced electrodes in the brain, as determined by histological screening after the completion of the behavioral experiments.

Fig 5 shows the positions of the electrode tips in the rat brain superimposed on a coronal section of a stereotaxic atlas map [11], where the MFB region is indicated by a grey area surrounded by a blue outline.

**Fig 5.**
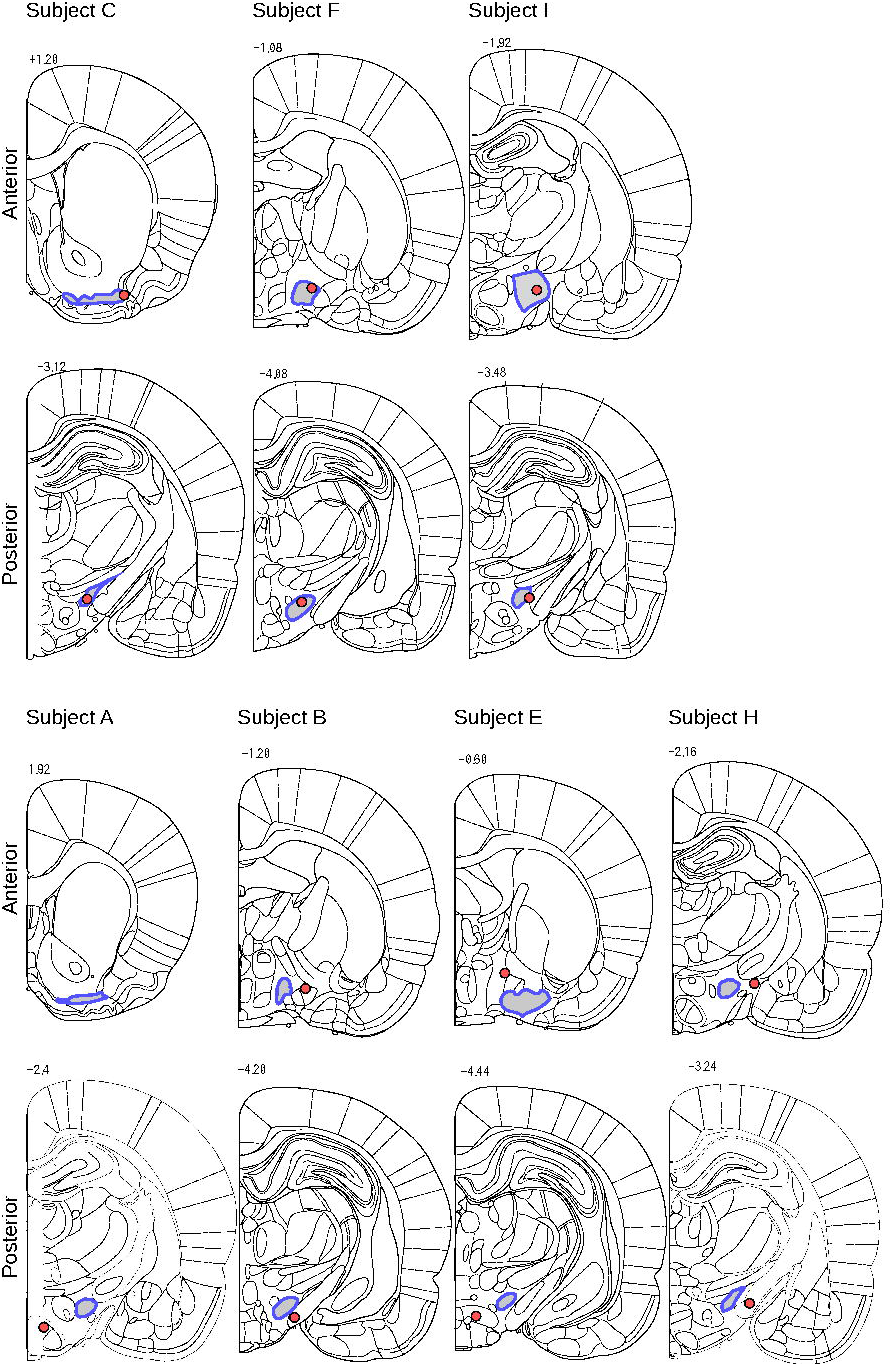
In each image, a red circle indicates the stimulated site and the gray area surrounded by a blue outline highlights the MFB region superimposed on a coronal section of a stereotaxic atlas image [11].

We found clear evidence that the electrode tips for both the anterior and posterior stimulation sites were located within the MFB in subjects C, F, and I, whereas the lesions were not located within the expected region in subjects A, B, E, and H. Both the anterior and posterior stimulation sites were located far from the MFB in subjects B, E, and H, respectively. In subject A, only the posterior site was located outside the MFB and the anterior site could not be determined in the observed sections.

Consequently, we categorized subjects C, F, and I in the *positive* group and subjects A, B, E, and H in the *negative* group, which were only tested on POW 1.

### Typical behaviors

Fig 6 shows typical rat locomotion trajectories. The rat moved all over the field, mostly along its walls when no electrical stimulation was provided.

**Fig 6.**
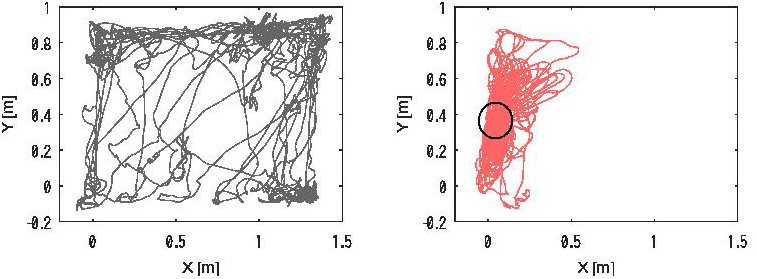
Typical rat trajectory without electrical stimulation in the brain (null condition; left), and a trajectory under stimulation in the MFB as the rat stayed in the virtual cage indicated by a circle (right).

In contrast, when we stimulated the anterior site in the MFB region as the subject entered the virtual cage, indicated by the black circle on the right plot, it repeatedly entered the virtual cage, mostly staying around the area.

Similarly, when we stimulated the posterior site in the MFB region of the subject as it entered the virtual cage, it produced the same behavior as above, remaining around the virtual cage.

### Behavioral analysis

This section presents the distance metric *d* value. In Figs. 7 and 8, the temporal progression of distance metric *d* through the trials is shown as a line plot.

**Fig 7.**
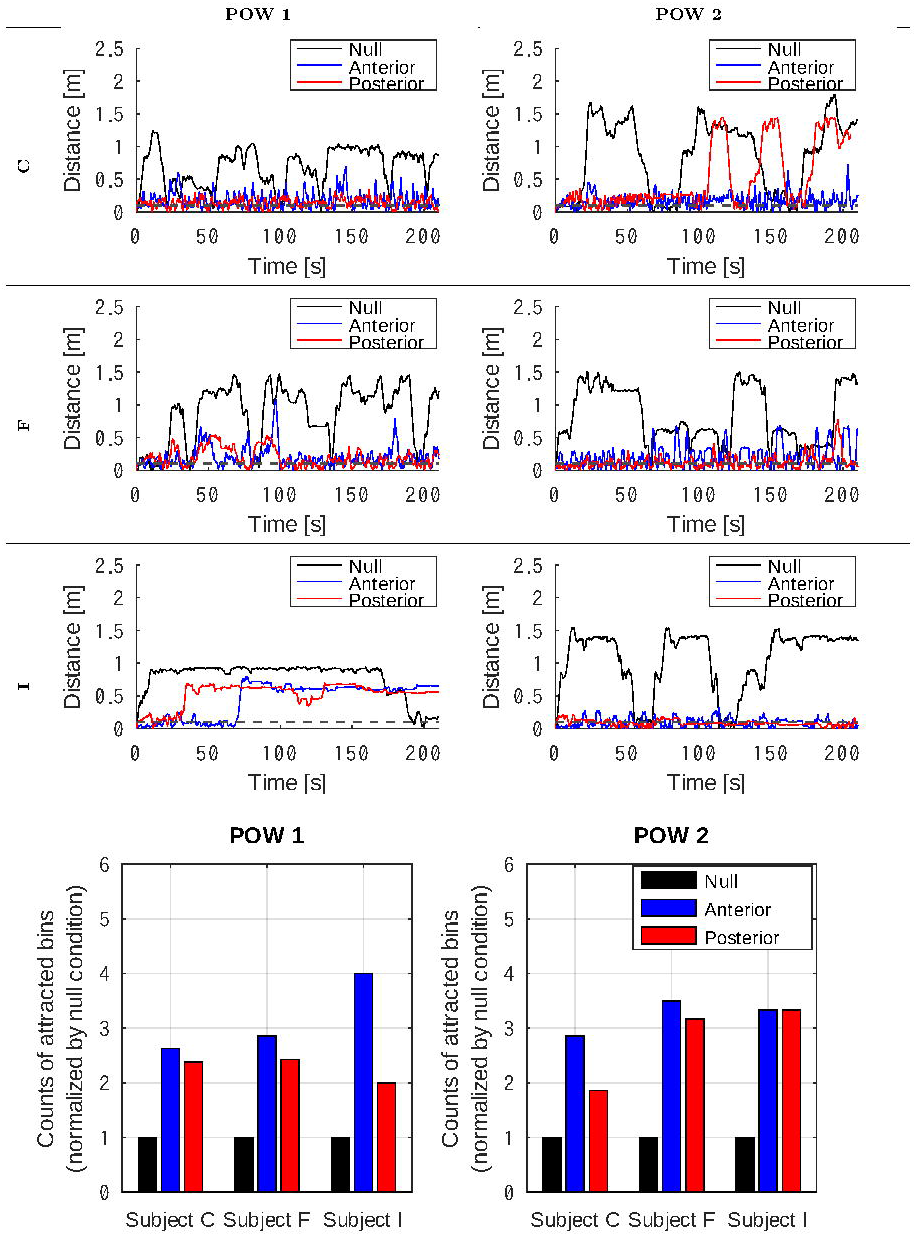
Line plots showing the temporal changes of the distance metric d throughout the sessions. The bar plots underneath compare the count of bins where the distance metric *d* stayed < 0.1 [m]. The analyses were conducted on POW 1 and 2 sessions for the subjects C, F, and I.

**Fig 8.**
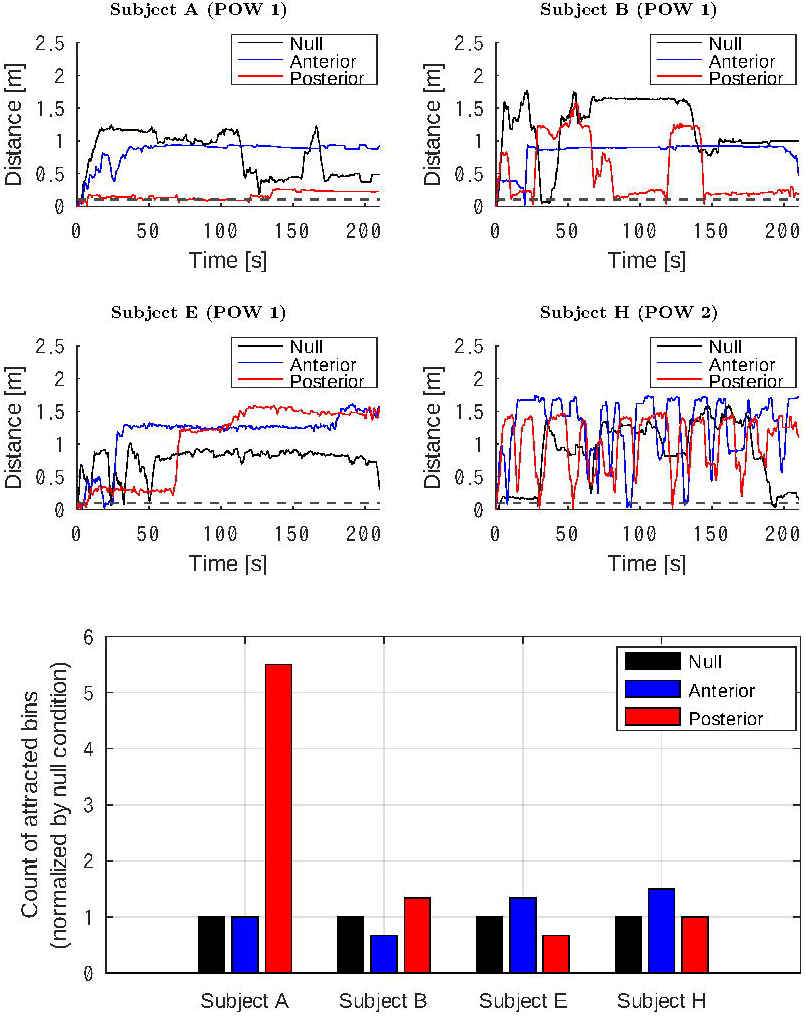
Line plots showing temporal changes in the distance metric *d* throughout sessions. Bar plots at the bottom comparing bin counts where the distance metric *d* was < 0.1 [m]. The analyses were conducted on subjects A, B, E, and H during the POW 1 session.

The bar plots compare the frame counts where *d* < 0.1 [m], that is, the rat was attracted to remain within the virtual cage. The counts during the anterior and posterior stimulation were normalized to those in the null condition.

#### Positive group

Line plots in Fig 7 show that subjects C and F stayed near the virtual cage throughout the sessions when stimulated at either the anterior or posterior site. The behavior of the rats was remarkably different from that in the null condition when they crossed the virtual cage. This preference was observed in POW sessions 1 and 2.

Subject I, on the other hand, left the virtual cage incidentally after staying around it for 30 – 70 s and failed to find the stimulation area in the POW 1 session.

Interestingly, it remained in the virtual cage during the POW 2 session.

The bar plots in Fig 7 indicate that all three subjects preferred staying in the virtual cage, and regardless of the stimulation sites and POW session, the counts of attracted frames increased with stimulation.

#### Negative group

The subjects in the negative group had no tendency to stay near the virtual cage despite the electrical stimulation of the brain. The line plots in Fig 8 indicate that the subjects and sessions, but not the stimuli, were responsible for changing the distance metric *d*.

Tge rats showed no remarkable difference in the counts of the attracted frames, except for subject A, which was less active in the POW 1 session and incidentally stayed near the virtual cage when the anterior site was stimulated. Although it yielded the highest number of attracted frames, the subject showed no response to the stimuli.

### Demonstration

Our Rattractor system showed the best performance for subject C during the first POW session. Here, we demonstrate how the subject was attracted to the virtual cage when it was either still or moving.

#### Still virtual cage

Fig 9 shows the number of times subject C stayed at a specific location divided by a 15 × 10 mesh.

**Fig 9.**
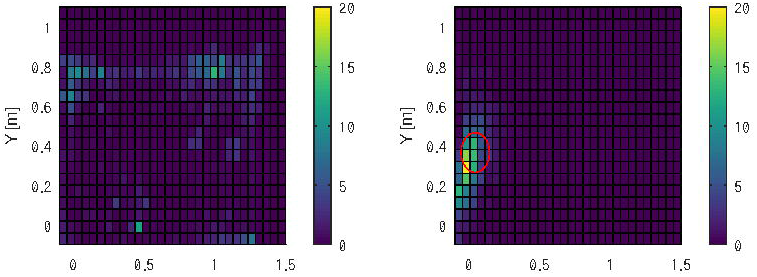
Histogram indicating the time s that the rat stayed on a specific location without stimulation (left), and in the virtual cage indicated by a red circle with stimuli in the MFB (right), respectively.

In the null condition without stimuli, the rats showed exploring behavior in the experimental field, yielding no remarkable focus points. In contrast, the rat preferred to stay near the virtual cage (illustrated by the red circle), where it could earn reward stimuli in the MFB.

The subject entered the virtual cage repeatedly, occasionally stepped out, and immediately returned to the area.

#### Moving virtual cage

Fig **??** shows the trajectory of subject C (light blue) and the virtual cage (dark red), which were moved manually. The virtual cage was initially set at the center of the field (X=0.75, Y=0.5), moved towards one corner (X=0.0, Y=1.0) at a constant velocity, and then moved towards the opposite corner (X=1.3, Y=0.0) before moving towards the approximate center of the field (X=0.75, Y=0.6).

The two plots below indicate the temporal changes in the x and y coordinates, demonstrating that the rat followed the virtual cage.

**Fig 10.**
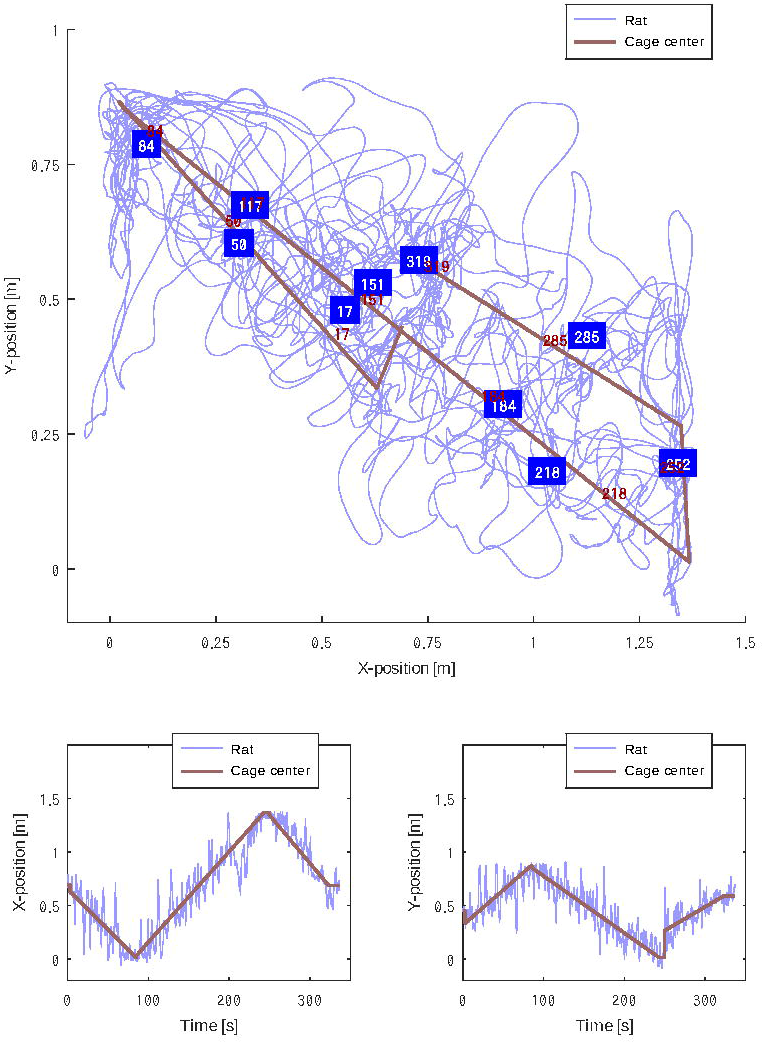
The plot above shows a two-dimensional trajectory of subject C (light blue) and the virtual cage (dark red), which was manually moved. The numbers show elapsed time s since the beginning of the session. The two plots below show temporal changes in x and y coordinates.

## Discussion & conclusion

This research aimed to provide a proof of concept of our “Rattractor” system, which can attract a rat into a virtual cage by providing electrical stimuli in the brain’s reward region. Three out of seven rats yielded successful results after placing a set of two needle electrodes in the MFB (Fig **??**), increasing the number of attracted bins compared to the null condition where no stimuli were applied (Fig 7).

Furthermore, subject C showed a clear preference towards staying near the virtual cage set at arbitrary locations (Fig 9), which was consistent when the virtual cage moved slowly (Fig **??**). Notably, the subjects followed the virtual cage even across the center of the experimental field, against their natural characteristics to avoid open spaces. A similar trend was observed in subjects F and I.

The performance of our system depended on electrode implantation. The electrodes of the three successful rats were correctly located in the MFB region according to subsequent histological analyses. In contrast, we did not find evidence of electrode tip locations within the MFB regions of the other four subjects, which showed null or avoidant responses to the stimuli.

Stimulation in the MFB region could drive the brain reward circuit by evoking dopamine-like responses in the striatum [12]. Moreover, it could cause enforced learning to gain more rewards, motivating the subject to search for and stay in the virtual cage. Thus, we aimed for a wide and long MFB region running parallel to the body axis to induce rewards effectively.

However, the correlation between the successful behavior and correct electrode placement indicates the relevance of electrode positioning in this system. We used a stereotaxic electrode manipulator to introduce the electrodes vertically into the brain, from the surface to the target regions. However, possible failure in the perpendicular grab of the electrodes or individual differences in skull/brain form and size may have resulted in failure to reach the target region. The actual electrode position could not be confirmed until the histological analysis was performed. Therefore, all behavioral experiments were conducted blind to the electrode positioning. Although we considered detecting the electrode tips using magnetic resonance (MR) imaging to avoid sacrificing the subjects, we concluded that the image accuracy would not be sufficient to determine the tip locations owing to potential artifacts.

Stimulation conditions, such as the current volume, frequency, and iteration number were determined according to a previous study [6] and adjusted to obtain the best performance. These parameters had to be strong enough to induce changes in locomotion but weak enough to avoid abnormal cell activity resulting in epileptic responses. According to the present results, both demands were satisfied by the system.

Rattractor is essentially a closed-loop behavior-reward feedback system that restricts animal behavior. Traditional approaches to restricting animal movement include the use of physical cages, walls, and choker rings. These are simple and effective but are not adaptable to various circumstances. Reward and punishment by a feeder allowed the animals to learn to stay within permitted areas. However, it requires training and is time consuming. By contrast, our reward system only requires the feeder to set a spatial region that can be set arbitrarily and adaptively, towards which the animal will be attracted. In our system, the animal was motivated to stay within the virtual cage without physically restricting its behavior, allowing it to resist the lead if necessary. Furthermore, the feeder could terminate rewarding at any time and free the animal from movement restrictions.

These characteristics are well-suited for grazing. For example, cattle and sheep could be led by the system within a farm without physical barriers, similar to shepherd dogs chasing sheep. Although implanting electrodes into all animals in a group is unrealistic, performing the surgery in a few leaders might be enough to guide the rest.

In conclusion, we developed a “Rattractor” system to automatically attract a rat towards a virtual cage only when the system was activated, allowing free movement when deactivated, which was only effective when the system was activated. We showed that our system could alter the behavior of a rat and guide it towards arbitrary locations if we succeeded in placing stimulation electrodes in the MFB. Developing a more precise surgical technique would increase the success rate of implants and would lead to the evaluation of the system with a larger number of subjects.

## Acknowledgments

This study was partially supported by JSPS Kakenhi for Young Researchers (B) 17K17684 and the Precise Measurement Technology Foundation (PMTP-F).

